# Gene Therapy for Cardiomyopathy associated with Duchenne Muscular Dystrophy in a Pig Model

**DOI:** 10.1101/2023.10.02.560452

**Authors:** Andrea Bähr, Petra Hoppmann, Tarik Bozoglu, Michael Stirm, Ina Luksch, Tilman Ziegler, Nadja Hornaschewitz, Samjhana Shrestha, Bachuki Shashikadze, Jan Stöckl, Nour Raad, Helmut Blum, Stefan Krebs, Thomas Fröhlich, Christine Baumgartner, Monika Nowak-Imialek, Maggie Walter, Christian Weber, Stefan Engelhardt, Alessandra Moretti, Nik Klymiuk, Wolfgang Wurst, Karl-Ludwig Laugwitz, Roger J. Hajjar, Eckhard Wolf, Christian Kupatt

## Abstract

**Background:** Genetic cardiomyopathies caused by mutations in the dystrophin gene *(DMD)* are only partially responsive to current pharmacological heart failure treatments, although dilated and arrhythomogenic phenotypes of cardiomyopathy are frequent.

**Objective:** In this study, we tested whether a normalization of Ca^2+^-handling by forced expression of SERCA2a in cardiomyocytes mitigates heart failure and arrhythmogenesis in a pig model for Duchenne muscular dystrophy (DMD).

**Methods and results:** Male offspring of pigs lacking *DMD* exon 52 are characterized by heart failure with reduced ejection fraction (HFrEF, EF 34.5±1.8% vs. 49.2±1.0% in control hearts), arrhythmogenesis due to large apical regions of reduced voltage amplitude and sudden cardiac death with a lifespan of usually less than 4 months. Slow antegrade intracoronary infusion of AAV1.SERCA2a (3×10^13^ virus genomes (vg) per pig) improved left ventricular ejection fraction (EF 47.3±2.0%, p<0.05) to a similar extent as germline editing of *DMD*Δ52 to *DMD*Δ51-52, inducing a Becker dystrophy (BMD) genotype (EF 46.7±3.8%). Moreover, AAV.SERCA2a significantly reduced myocardial inflammation and fibrosis and areas of reduced AP amplitude.

**Conclusions:** In *DMD* pigs, 3×10^13^vg/heart of GMP-grade AAV1.SERCA2a sufficed to normalize left ventricular function and improved electrical vulnerability of the heart. Hence, AAV.SERCA2a may serve as a treatment option for DMD cardiomyopathy.

## Introduction

Duchenne muscular dystrophy (DMD) is a devastating genetic disease caused by loss of dystrophin, a protein attaching the cytoskeleton to the cell membrane. Loss-of-function mutations of this x-chromosomal gene affect boys at a rate of 1:3500, constituting the most frequent form of muscle dystrophy (1,2). Subsequent to progressive skeletal muscle decay, patients loose ambulation in their first decade of life, experience dyspnoea due to thoracic and diaphragm weakness and develop - mostly in the second to third decade of life - a distinct form of cardiomyopathy. In contrast, the phenotype of Becker muscular dystrophy, which occurs at about 1/3 of the DMD rate (3), is attenuated by a shortened, but largely functional dystrophin. Here, symptoms vary from mild to more severe impairment of muscle function, although the majority of boys with Becker dystrophies also develop a distinct cardiomyopathy (1,4,5).

Beyond pharmaocological therapy - initiating ACE-inhibitors even before the onset of symptoms at teenage age (6), potentially extending towards mineralocorticoid antagonism (7) and full-blown heart failure therapy later on (8–10) - little specific therapies have been established for DMD cardiomyopathies despite the unmet clinical need. Lately, antisense oligonucleotides (AONs) have been conditionally approved for treatment of muscle weakness (11), with moderate increases of dystrophin expression in the skeletal muscle, and their impact for heart function largely unknown. A more effective approach towards restoring dystrophin is currently under intense investigation in phase-3 trials: testing adeno-associated viruses (AAVs) encoding microdystrophins for ubiquitous muscular expression, attempting to convert a Duchenne to a Becker phenotype. However, besides high AAV-doses bordering at toxic side effects, the impact on hearts is known only from preclinical experiments in *mdx* mice (12,13), which rarely develop a severe phenotype with or without therapy.

Alternately, correction of the disease-causing mutation has been achieved via Cas9-mediated editing of the DMD gene in preclinical models of mice (14–17), dogs (18) and pigs (19). In each successful case, a gain of muscle strength or heart function was detected after re-expression of dystrophin, though in a shortened form lacking the mutated and, if appropriate, the neighboring exons. However, cardiomyocyte editing may raise concerns with respect to safety and efficacy, since current protocols of cardiac gene editing require high doses of AAVs encoding active Cas nucleases and achieve only incomplete gene editing of the heart. Representing the optimal outcome of such a strategy, we performed germline gene editing of dystrophin (identical to the approach used in adult pigs, cf. 19), generating a Becker muscular dystrophy (BMD) pig model, which we include in this study.

However, till the arrival of efficient and safe gene editors for DMD, a more feasible and applicable gene therapy approach may serve the current need of patients suffering from DMD heart failure Therefore, in the current study, we tested whether strengthening of the sarcoplasmatic reticulum (SR) Ca2+ ATPase SERCA2a, a calcium pump lacking in end-stage heart failure, would ameliorate heart function in DMD pigs. Of note, Duan et al. were able to successfully treat mdx mice with a one-time AAV.SERCA2a vector, restoring skeletal muscle and heart function for a period of ∼2 years (20). Focusing on DMD-cardiomyopathy, we loco-regionally transduced DMD-hearts with AAV1.SERCA2a via coronary injection of the LAD and RCx and injection through a microcatheter. Three weeks later, AAV-SERCA2a was expressed in the heart tissue distal to the injection site and imrpoved function and structure of the DMD hearts in a broad manner, normalizigng not only systolic funcion, but also microvascular density and longterm inflammation of the heart.

## Methods

Pigs were purchased from Department of Veterinary Medicine, LMU Munich (Oberschleißheim, Germany), Animal care and all experimental procedures were performed in strict accordance to the German and National Institutes of Health animal legislation guidelines and were approved by the Bavarian Animal Care and Use Committee.

### Pig generation

A heterozygous DMDΔ52 mutation was introduced in female primary kidney cells (German Landrace × Swabian-Hall hybrid background) by homologous recombination with a modified bacterial artificial chromosome (BAC) in which DMD exon 52 was replaced by a neomycin resistance cassette (Fig. 1A). The loss of exon 52 causes a shift of the reading frame, resulting in premature stop codons (Fig. 1B). A correctly targeted single-cell clone was used for somatic cell nuclear transfer (SCNT). Transfer of cloned DMD+/- embryos to recipients resulted in the birth of 7 live DMD+/- piglets, of which one (Fig. 1C) could be raised to adulthood and served as founder of the breeding colony of DMD pigs used in the present study.(21)

**Figure 1:**
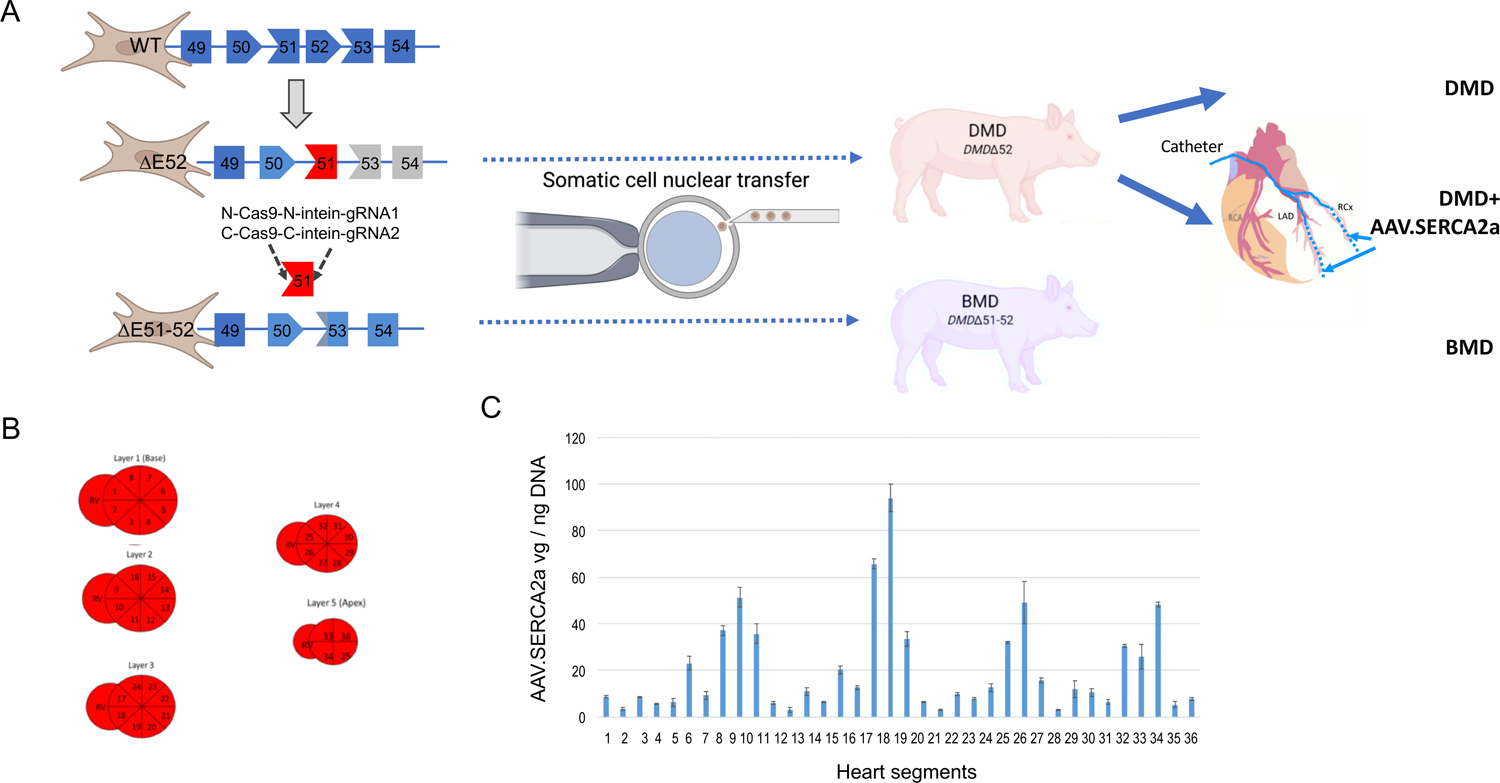
Models of DMD and BMD pigs. **A** Wildtype (WT) porcine fibroblasts were subjected to elimination of *DMD* exon 52 by homologous recombination. Somatic cell nuclear transfer (SCNT) of fibroblast nuclei lacking *DMD* exon 52 enabled generation of a pig cohort with absent dystrophin expression (21). Additional Cas9-mediated excision of *DMD* exon 51 was performed in somatic porcine cells of this genotype, resulting in expression of a shortened, but stable *DMD*Δ51-52. Via a second round of SCNT, Becker muscle dystrophy (BMD) pigs were generated, which were used in this study (23). **B** *In vivo* transduction was conducted by antegrade intracoronary infusion of AAV1.SERCA2a into the LAD and RCx under nitroglycerin co-infusion (cf. Methods, (). **C** Systematic probing for AAV transduction yielded up to 90 AAV genomes /ng DNA. Representative example of 3 experiments.

For the generation of DMDΔ51-52 (BMD) pigs, DMD exon 51 was deleted in a primary kidney cell line (PKC) of a male DMDΔ52 pig, as described before (19). Briefly, porcine fibroblasts were co-transfected with two plasmids, each carrying the sequence for a guide RNA (gRNA) binding upstream (AGAGTTCCTAAGGTAGAGAGAGG) or downstream (ATAAAGATAAGAGCTGGCAGAGG) of exon 51 as well as the sequences of an intein-mediated split-Cas9 system (21). After a 24-h resting period, clonal separation was performed. Single-cell clones were picked and screened for the deletion of exon 51 by PCR. The resulting PCR products were verified by gel electrophoresis and Sanger sequencing. *DMDΔ51-52* cell clones were used for somatic cell nuclear transfer (SCNT)(42). Offspring were genotyped by PCR detecting the deletion of DMD exon 51.

### AAV1.SERCA2a generation

The GMP clinical grade AAV1.SERCA2a vector was produced as described previously by Jessup et (22). The AAV1 viral particle was constructed as a hybrid or pseudotype vector in which the vector genome incorporates the capsid sequence from AAV1 and the inverted terminal repeat from native AAV2. In conjunction, the vector incorporates the human SERCA2a expression cassette under the control of the cytomegalovirus enhancer/promoter.

### Regional AAV1.SERCA2a application

DMD pigs were anesthetized and instrumented as previously described (23–25). In the current study, we regionally applied AAV1.SERCA2a at a dose of 3×10^13^ virus genomes (vgs) via antegrade application at d0: the vector-containing solution was infused into the left anterior descending artery (LAD, 60%) and into the circumflex artery (RCx, 40%), each using a ballon (2.5×12mm) inflated at 4 bar for 3min to slow blood flow into the target region and to increase contact time for the drug infusion (Fig.1A). 0.2mg nitroglycerin were infused to facilitate transendothelial passage. 2 pigs weighing < 15kg at an age of 2.5months were excluded from the study, for an increased risk of vessel injury during instrumentation due to small size, and a high risk of sudden cardiac death while waiting for further growth. No animal was lost subsequent to instrumentation and AAV1.SERCA2a application.

### AAV1.SERCA2a detection

DNA was extracted from 50µm cryostat sections of left ventricular tissue samples with NucleoMag Tissue Kit (Macherey-Nagel, Germany) as per manufacturer’s instructions. AAV genome copy numbers were quantified with qPCR using FastStart Probe master mix (Sigma, Germany) on a CFX96 thermal cycler (Bio-Rad, Germany). The primer – hydrolysis probe set used for quantification was directed against the coding sequence of SERCA2a transgene (F: 5’ AAC CCT CCC ACA AGT CTA AAA TC 3’, P: 5’ AGC ATC GTT CAC GCC A 3’, R: 5’ CTC GGC TTT CTT CAG AGC AG 3’).

### Proteomics of left ventricle tissue

Sample preparation for proteomics of cardiac tissue was performed as described recently (26). Briefly, snap frozen tissue was pulverized, and the resulting powder was lysed in urea 8 M/50 mM ammoniumbicarbonate using sonication. Subsequent to reduction and alkylation of cysteine residues, samples were digested with Lys-C for 4 h followed by trypsin for 16 h at 37 °C. For MS-based proteomics, 1 µg peptides were injected on an Ultimate 3500 RSLC coupled to a Q-Exactive HF-X (Thermo Fisher Scientific). Peptides were first trapped using a PepMap 100 C18 trap column and then separated with a PepMap C18 analytical column (75 µm x 50 cm; 2 µm; both Thermo Fisher Scientific) at 250 nL/min with a two-step gradient: first from 3% B to 25% B in 30 min, then to 40% B in 5 min. Eluent A was 0.1% formic acid in water and B was 0.1% formic acid in acetonitrile. MS spectra were acquired in data dependent mode with 12 product spectra per cycle.

Raw data were processed with MaxQuant (1.6.7.0)(27) using all NCBI RefSeq protein *Sus scrofa* entries (downloaded in July 2020). For quantification, a label free acquisition approach was chosen. The false discovery rate was controlled to be 1%. Data analysis was done with Perseus(28), R(29), the R packages tidyverse (https://tidyr.tidyverse.org) and ComplexHeat-map (30). The mass spectrometry proteomics data have been deposited to the ProteomeXchange Consortium via the PRIDE partner repository with the dataset identifier PXD045592 (https://www.ebi.ac.uk/pride/).

### Hemodynamic measurements

Pigs were anesthetized and instrumented as previously described (23–25). Briefly, global myocardial function was assessed by pressure-tip catheter placement in the left ventricle (for LV enddiastolic and systolic pressures) at rest and rapid atrial pacing (150/min), whereas analysis of ejection fraction was performed after LV angiography in anterior-posterior position (yielding slightly smaller control values than a right anterior oblique view).

### Electrophysiological assessment

The Rhythmia mapping system was used for high-resolution 3D-mapping (Boston Scientific, Natick, Massachusetts), as described before (19). Briefly, bipolar activation maps were created in wildtype hearts, untreated DMD and BMD hearts, and SERCA2a-treated hearts. An I8.5F IntellaMap-Orion catheter (Boston Scientific, USA) containing 64 flat microelectrodes (0.8 mm diameter) in a basket configuration with 8 splines can be opened and closed to provide appropriate wall contact for detection of electrophysiological signals. The LV surface geometry was generated by including all points recorded within 2 mm from the outermost surface of the map (defined by outermost reach of any of the electrodes in 3D space). The voltage for bipolar electrograms was derived measuring from peak to peak. The low-voltage area and endocardial scar are were defined on the bipolar voltage map as <1.3mV and <0.3 mV, respectively.

For quantitative analysis of the electroanatomical maps, maximal, minimal and mean bipolar electrogram voltage was calculated for each LV-map. Quantification of low voltage scar areas - defined as bipolar voltage <1.3mV – was done using the paraView open-source, multi-platform data analysis and visualization application (Kitware, Clifton Park, New York, USA).

### Fibrosis staining

Cryosections (8 µm) were generated using standard histological techniques. For quantification of collagen deposition, myocardial sections were stained with Sirius red and Fast green. Images were acquired using a Leica Thunder microscope (Leica, Hilden, Germany) with a 10x objective and fibrosis was quantified using Image J software (NIH). After random sampling, 10 cryosections were analyzed per pig.

### Microvascular density

For pig immunofluorescence, heart samples were frozen on dry ice, embedded in OCT and sectioned on to slides. Heart slides were fixed in cold Acetone, washed in PBS (5 min. 3 washes) and blocked with 10% NFS, for 60 min. Then, slides were incubated overnight at 4°C with the following antibodies (diluted in blocking buffer): Anti PECAM-1 (sc-376764, Santa Cruz, 1:50), anti NG2 (NG2 antibody, AB5320 Millipore, Massachusetts, USA, 1:200). The slides were washed with PBS (5 min,3 washes), stained with the relevant secondary antibody for 2 hours at RT. Slides were mounted and visualized. 10 slides were analyzed per pig.

### CD45 Staining and quantification

Heart slides were acetone-fixed and blocked with goat serum, followed by overnight staining with pig specific CD45 antibody (clone K252.1E4, Bio-Rad, 1:200). After secondary ab and DAPI applications, slides were mounted and imaged with a Thunder Imaging System (Leica, Germany). Entire sections were imaged at 200X magnification and stitched together. CD45 signal was thresholded (50–255) using ImageJ and quantified as integrated density percent of total area. 10 random samples were analyzed per pig.

### Quantitative real-time PCR

Total RNA was extracted from tissue using and reverse-transcribed into cDNA. qRT-PCR was performed on an iQ-cycler (Bio-Rad, Munich, Germany) with SYBR-Green Supermix (Bio-Rad, Munich, Germany). For qPCR, the following primers were used: *GAPDH* forward: 5’-AATTCAACGGCACAGTCAAG-3’, reverse 5’-ATGGTGGT-GAAGACACCAGT-3’; *NRF2* forward: 5’-GGGGTAAGAATAAAGTGGCTGCTC-3’, reverse: 5’-ACATTGCCATCTCTTGTTTGCTG-3’

## Results

As depicted in Fig. 1A, porcine models of Duchenne (*DMDΔ52*) and Becker (*DMDΔ51-52*) muscular dystrophy were generated by somatic cell nuclear transfer (s. Methods). Animals of the Duchenne cohort were treated by intracoronary artery infusion of AAVs containing SERCA2a for treatment of the DMD cardiomyopathy. After termination of the experiments and systematic division of the heart into 36 segments (Fig. 1B), AAV.SERCA2a was found in all analyzed myocardial segments, with a preference to the well-muscularized septal and anterior regions (Fig. 1C).

Although we transduced a single gene sequence into cardiomyocytes, the likelyhood of altering expression patterns in more profound ways seemed high, since DMD itself causes such alterations (19,26). Principal component analysis of the proteome confirmed that the global protein profile of DMD hearts, which differed considerably from WT hearts, was partially rescued in SERCA2a-treated DMD hearts (Fig. 2A). Interestingly, significantly dysregulated proteins detected in DMD hearts showed a trend towards normalization in the SERCA2a-treated DMD-hearts (Fig.2B). Pro-inflammatory acute phase proteins such as inter-alpha-trypsin inhibitor heavy chain H4 (ITIH4) or alpha-2-macroglobulin (A2M), elevated in the instance of DMD, were downregulated upon AAV.SERCA2a, whereas the opposite was found for the mitochondrial protein delta(3,5)-delta(2,4)-dienoyl-CoA isomerase (ECH1) (Fig.2C-E). The cardiomyocyte-specific sarcomere gene myosin heavy chain 6 (MYH6), the most downregulated proteins in DMD-hearts, was significantly increased upon treatment (Fig. 2F). Next, we analyzed whether these profund changes in the proteome translated into structural improvements of the DMD hearts, which are known for their fibrotic alterations. Indeed, in histological analysis, the fibrotic area of the myocardium was significantly increased in untreated DMD compared to WT. Of note, this was not the case in BMD animals. Remarkably, SERCA2a-treatment of DMD hearts ameliorated interstitial fibrosis (Fig.3A). Moreover, microvascular rarifaction, a hallmark of malperfusion of failing muscle in DMD (19), was present in DMD hearts, but largely improved in both, BMD corrected and SERCA2a-treated DMD hearts (Fig. 3B). Inflammation, a key driver of skeletal muscle decay in skeletal muscle, which is regularly treated by steroids in DMD patients, is also present in untreated myocardium, as depicted by recruitment of CD45^+^ cells (Fig.3C). Of note, SERCA2a-treatment reduced this process to a much lower level, though still elevated when compared to control and BMD groups (Fig. 3C).

**Figure 2:**
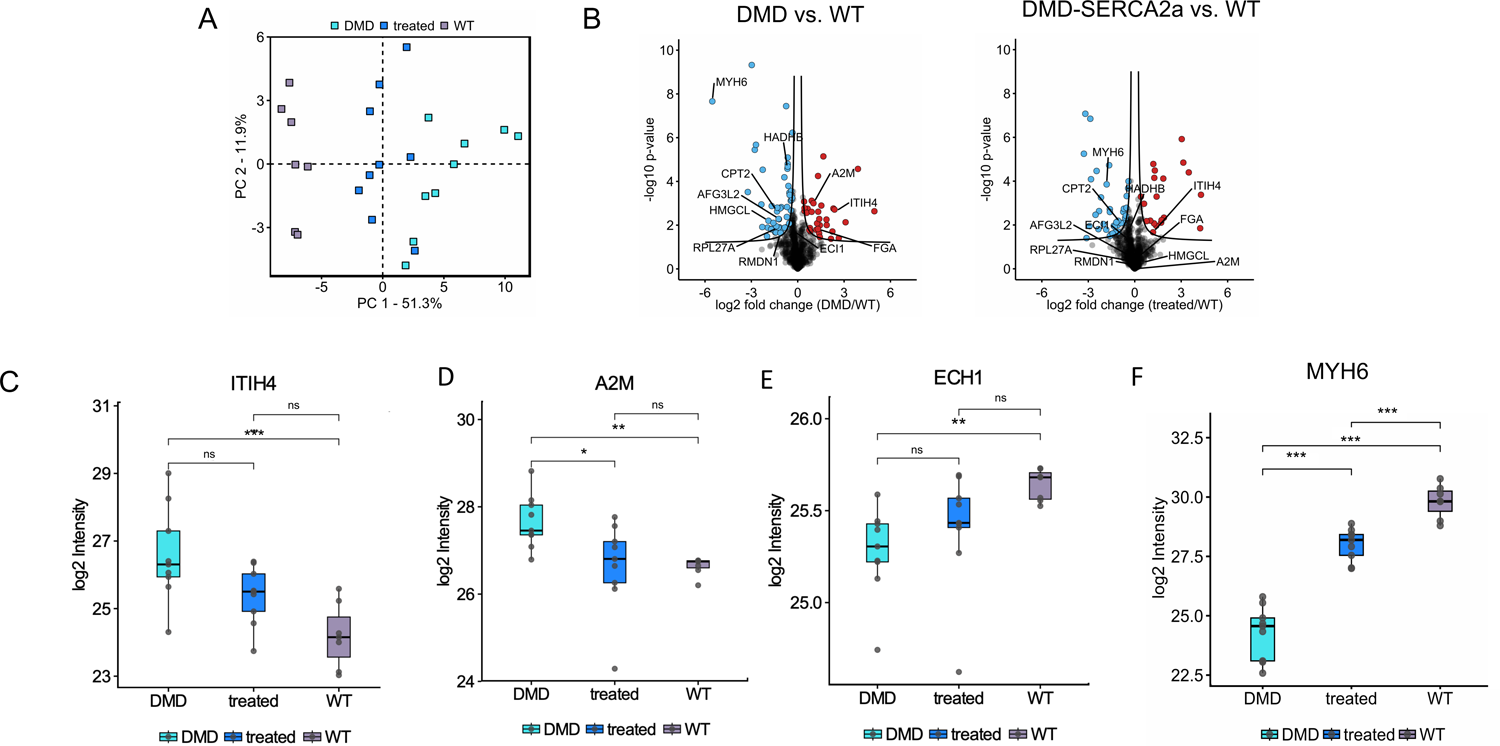
Proteomic profile of DMD and DMD+AAV.SERCA2a. **A** Principal component analysis (PCA) reveals the capability of AAV1.SERCA2a (blue squares) to shift the DMD proteome (turquoise squares) towards WT (grey squares), though congruence was not achieved. **B** Volcano blots of proteins differently expressed in DMD vs WT and DMD-SERCA2a vs. WT reveal that selected proteins are shifted towards normalcy by AAV.SERCA2a therapy, though not reaching WT levels. **C** Protein levels of MYH6 as one of the most downregulated proteins in DMD, increasing upon AAV.SERCA2a treatment. D The acute phase protein inter-alpha-trypsin inhibitor heavy chain H4 (ITIH4) and alpha-2-macroglobulin (A2M) were downregulated by AAV.SERCA2a, whereas the mitochondrial protein delta(3,5)-delta(2,4)-dienoyl-CoA isomerase (ECH1), downregulated in DMD, was found increased after AAV.SERCA2a application.

**Figure 3:**
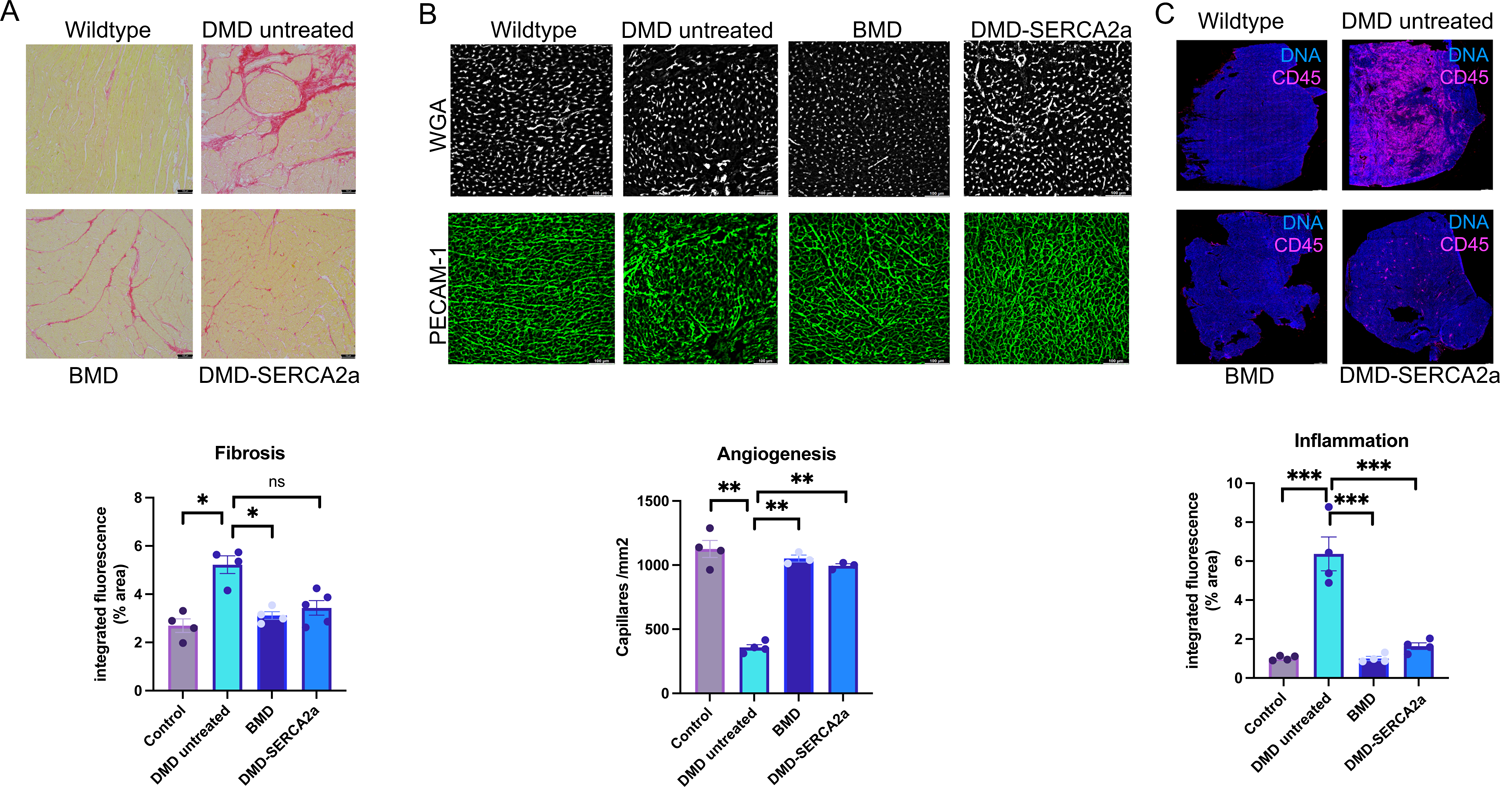
AAV.SERCA2a augments microcirculatory density. **A** Examples of interstitial fibrosis in DMD, BMD and DMD+AAV.SERCA2a. Quantification reveals no difference among experimental groups. **B** Examples and quantification of capillary density, assessed by PECAM-1 staining of endothelial cells, did not reveal a difference between WT and BMD hearts, whereas DMD hearts displayed significantly lower levels of microvascular density. AAV.SERCA2a, however, increased PECAM-1^+^ cell density. **C** Low magnification microscopy revealed an increased amount of CD45^+^ cells recruited to LV myocardium, compared to WT and BMD hearts. AAV.SERCA2a treatment reduced the CD45^+^ leukocyte level. Data are mean ± SE, P values were determined by unpaired Mann-Whitney U test.

Since the main stigmata of failing hearts, namely fibrosis, capillary rarefaction and inflammation, were improved by AAV1.SERCA2a, we probed the hypothesis that cardiac function would benefit from this molecular interention. Intracoronary AAV1.SERCA2a-infusion improved the ejection fraction of DMD pigs, indistinguishable to the level of wildtype pigs (Fig. 4A) and BMD pigs. Accompanying hemodynamic data indicated a trend towards improved contraction velocity compared to untreated DMD animals, similar to BMD animals (Fig. 4B). The LVEDP measurements indicated a decrease from a heightened untreated DMD-level (28.3±0.9mmHg) towards normalcy (16.5±0.8mmHg).

**Figure 4:**
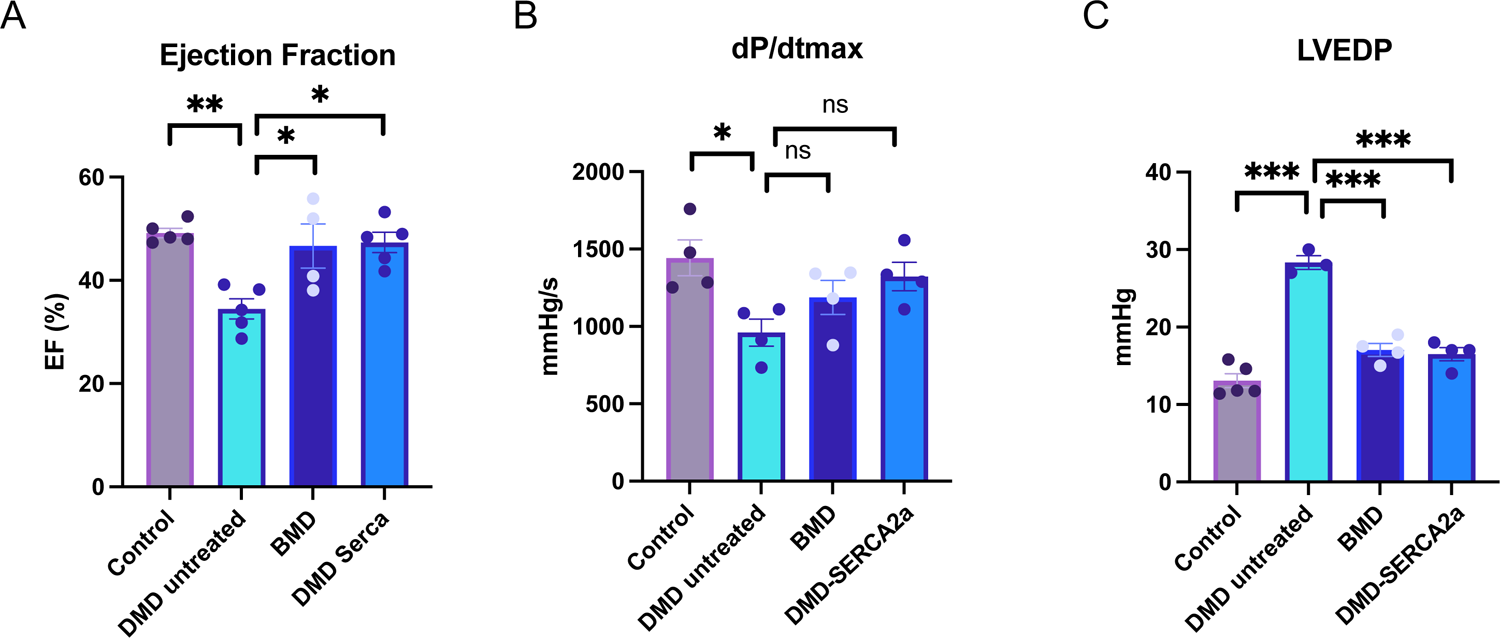
Functional impairment of DMD hearts are attenuated by AAV.SERCA2a. **A** Ejection fraction (EF) and **B** contraction velocity (dP/dt_max_) were found decreased in DMD, unless AAV.SERCA2a was applied. application improved systolic function significantly. **C** Left ventricular enddiastolic pressure (LVEDP) levels were found increased in DMD, but returned to WT levels after AAV.SERCA2a treatment. Data are mean ± SE, P values were determined by unpaired Mann-Whitney U test.

After documenting the improved LV function in AAV1.SERCA2a treated DMD animals, we analyzed the impact of SERCA2a gene therapy on voltage amplitude in electrophysiologic analysis. In Fig.5, high resolution maps (Rhythmia, Boston, USA) indicate that DMD hearts carry large areas of no to low amplitude, mostly in the mid-to-apical part of the ventricle, matching the structural correlate of increased fibrosis. BMD completely normalized this electrophysiological alteration. In SERCA2a-treated animals, a overall reduction of low-amplitude areas was quantified (Fig.5B).

**Figure 5:**
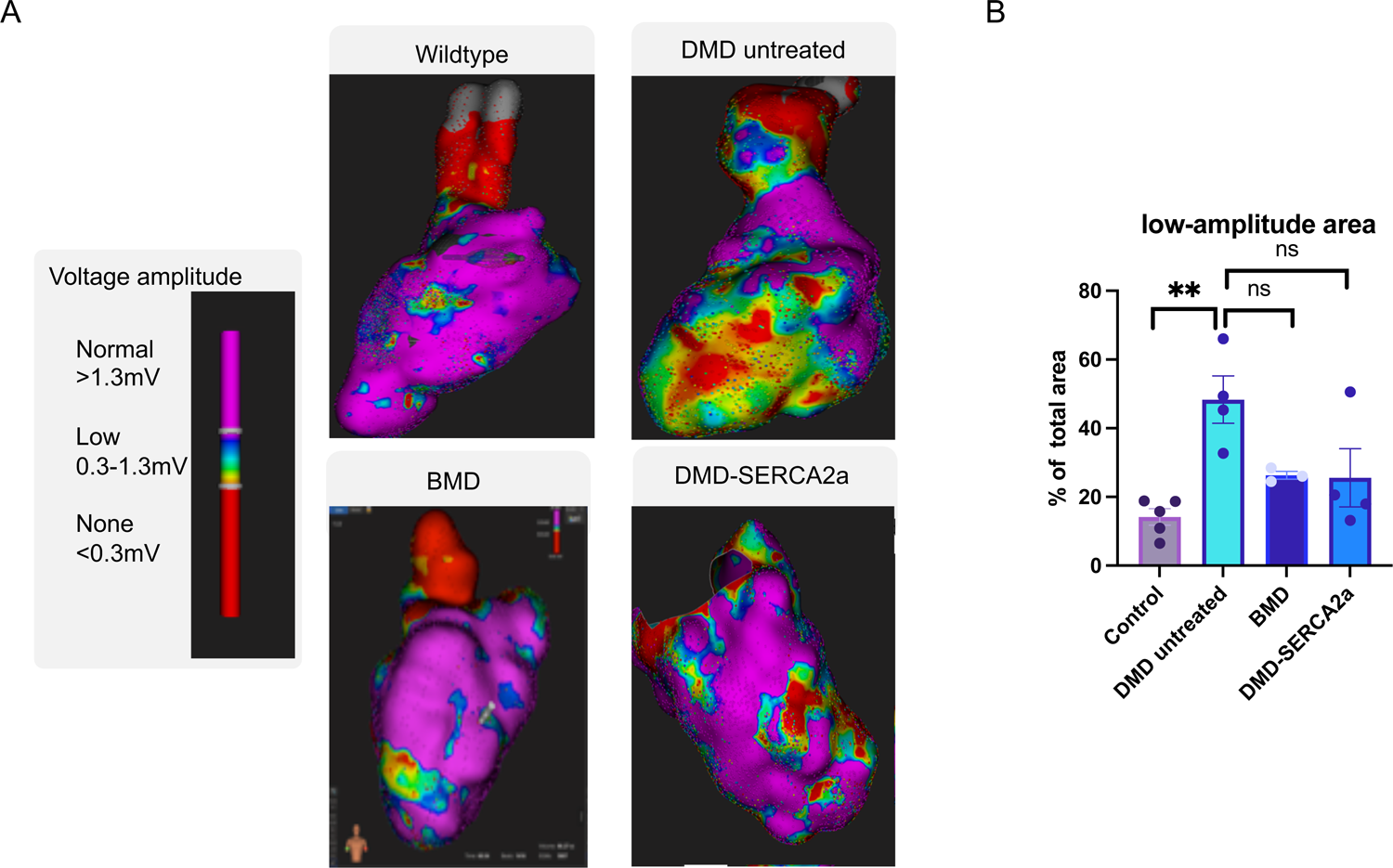
Electrophysiologic mapping of DMD&AAV.SERCA2a. **A** Examples of high-resolution electrophysiologic maps of wildtype, DMD, BMD and DMD hearts, with or without AAV.SERCA2a transduction. **B** Quantification of low- and no-voltage amplitude areas revealed that DMD-hearts contained significantly larger areas of low- and no-amplitude, as opposed to BMD pigs. AAV.SERCA2a treatment revealed an intermediate phenotype. Mean ±SEM, *=p<0.005 vs. Control. #=p<0.05 vs. Control

## Discussion

Our study establishes the potential of locoregional expression of SERCA2a through AAV1 application to improve heart function in DMD: a moderate dose of 3×10^13^ vg/heart, infused into the left coronary arteries (LAD and RCx, cf. Fig.1), tended to normalize a broad spectrum of proteins (Fig.2), improved cardiac fibrosis, microcirculatory density and chronic inflammation (Fig. 3). Functionally, intracoronary AAV1.SERCA2a treatment enhanced ejection fraction and contraction velocity (Fig.4), to similar levels to the ones obtained from ubiquitous gene editing of dystrophin in BMD pigs. Moreover, electrophysiological mapping revealed that AAV1.SERCA2a-treated DMD hearts in aggregate displayed less areas of low or no voltage (Fig.5).

Heart failure with reduced ejection fraction is a disease state caused by a wide variety of pathophysiologic entities such as ischemic, inflammatory, structural diseases as well as genetic cardiomyopathies, including DMD. In the latter case, despite minimal physical demand due to an early loss of ambulation, heart failure and arrhythmias become apparent from the second to third decades of life on. In fact, since aberrant calcium handling is a constitutive element of ionic imbalances common in late stage heart failure, additional expression of SERCA2a is an attractive treatment option. In DMD, preclinical murine *mdx*-models have revealed a role of sarcolipin as a molecular brake of calcium storage by Serca (31–33), and an improved muscle strengh with pharmacologic activation of Serca (34).

These observations are consistent with a previous report of AAV9.SERCA2a in *mdx*-mice, where a single systemic administration of 6×10^12^ vgs of AAV9.SERCA2a at 3 months of age sufficed to improve heart function with regard to ejection fraction and pressure volume loops at 21 months (20). Of interest, despite the continuous absence of dystrophin, cardiac fibrosis was also attenuated after AAV9.SERCA2a treatment, limiting the presence of arrhythmogenic substrates, similar to our study.

These findings, among others, point to a broad spectrum of alterations induced by DMD, of which activation of fibrosis and inflammation are the most consistent ones (Fig.3) (35), implying acute phase proteins such as alpha-2-HS-glycoprotein (AHSG), alpha-1-acid glycoprotein 1 (ORM1), inter-alpha-trypsin inhibitor heavy chain H4 (ITIH4) and alpha-2-macroglobulin (A2M), the latter two tending towards normalcy after AAV.SERCA2a application (Fig. 2C,D). Moreover, the downregulation of mitochondrial proteins, such as, 2,4-dienoyl-CoA reduc-tase (DECR1), enoyl-CoA delta isomerase 1 (ECI1) and delta(3,5)-delta(2,4)-dienoyl-CoA isomerase (ECH1), the latter upregulated upon AAV.SERCA2a (Fig. 2E), hint at mitochondrial dysfunction, whereas downregulation of proteins from the mitochondrial membrane and matrix (e.g., glutaryl-CoA dehydrogenase [GCDH], cytochrome b5 type B [CYB5B], and sirtuin 3) indicates mitochondrial instability. Additionally, ribosomal genes providing transcription and proteins of the ß-dystroglycan complex are less expressed, pointing at a downregulation of transcription as well as plasmalemma stability and signaling (35).

In this regard, AAV1.SERCA2a shifts some of the dysregulated proteins towards a wildtype level, although not achieving congruence at the timepoint measured (3 weeks after AAV1.SERCA2a application, cf. Fig2), as exemplified by the strongly downregulated sarcomere gene *MYH6* that shifted towards normalcy after AAV1.SERCA2A application (Fig.2F).

Moreover, consistent with the findings in AAV9-SERCA2a treated mice, CD45^+^-leukocyte recruitment is downregulated in AAV1.SERCA2a treated myocardium, when compared to untreated DMD hearts (Fig.3C). In addition, capillary rarefaction, as detected in the affected DMD muscles, appears to accelerate muscle dysfunction and is sensitive to treatment with pro-angiogenic factors (36–38). Consistent with an improved systolic function upon increased SERCA2a expression, capillary density was found increased in SERCA2a-treated muscle (Fig.3B).

As demonstrated recently, myofibroblast activation and collagen deposition make muscle and heart function declining over time, a fate at least in part attenuated by antifibroblast microRNA therapies (39,40). In this regard, a reduction of cardiac fibrosis was detected 3 weeks after AAV.SERCA2a treatment in DMD hearts (Fig. 3A). The high-resolution electrophysiologic mapping, though, indicated a decrease in low-amplitude areas. These observations point to a normalization of ion imbalances in the cardiomyocyte and a decreased risk of arrhythmogenicity. Na_v_1.5, which colocalizes with dystrophin and syntrophin (41) and has been described as downregulated in *mdx* mice(42), has not been found to be upregulated by AAV1.SERCA2a in this and a previous study (43). In a report by Lyon et al (44) SERCA2a gene therapy stabilized sarcoplasmic reticulum (SR) Ca load, reduced ryanodine receptor phosphorylation and decreased SR Ca leak, reducing ventricular triggered activity in vitro and in vivo. Normalization of intracellular Ca handling (19) most likely contributes to the salutary effects of AAV1.SERCA2a on electrophysiological parameters in DMD pigs.

### Study limitations

In this study, we deliberately chose a dose of 3×10^13^ vgs of AAV1.SERCA2a for locoregional transduction by intracoronary application. Though it is conceivable that other applications such as retrograde infusion could achieve similar or even better results, this route has been used clinically before and appears as safe and practical.

## Conclusion

In this study, we demonstrated the feasibility of restoring systolic function in a preclinical porcine model of genetic DMD cardiomyopathy for the first time. Far beyond its own overexpression, SERCA2a treatment partially normalized a variety of dysregulated proteins, decreasing arrhythmogenecity as well as inflammation and capillary rarefaction. These results pave the way to clinical application of AAV1.SERCA2a in DMD patients, which is currently under way for adult patients with DMD induced cardiomyopathy.

## Acknowledgements

We thank Anja Wolf for expert technical assistance.

